# ABCA1 deficiency causes tissue-specific dysregulation of the SREBP2 pathway in mice

**DOI:** 10.1101/2024.02.20.580966

**Authors:** Yoshio Yamauchi, Sumiko Abe-Dohmae, Noriyuki Iwamoto, Ryuichiro Sato, Shinji Yokoyama

**Affiliations:** Department of Applied Biological Chemistry, Graduate School of Agricultural and Life Sciences, The University of Tokyo, Bunkyo-ku, Tokyo 113-8657, Japan; Department of Biochemistry II, Nagoya University Graduate School of Medicine, Nagoya 466-8550, Japan; Biochemistry, Nagoya City University Graduate School of Medical Sciences, Nagoya 467-8601, Japan; Department of Food and Nutritional Sciences, Chubu University, Kasugai 487-8501, Japan

**Keywords:** ABCA1, HDL, SREBP2, Tangier disease

## Abstract

The ATP-binding cassette transporter ABCA1 plays an essential role in the formation of high-density lipoprotein (HDL) by mediating phospholipid and cholesterol efflux to apolipoprotein A-I. In humans, loss-of-function mutations in the *ABCA1* gene cause Tangier disease (TD), a familial HDL deficiency. In addition to the disappearance of HDL, TD patients and *Abca1*^-/-^ mice exhibit the cholesterol deposition in peripheral tissues through a mechanism poorly understood, which may contribute to the development of premature atherosclerosis. We and others have shown that ABCA1 deficiency causes hyperactivation of the sterol regulatory element-binding protein 2 (SREBP2) pathway *in vitro*. In this work, we investigate whether ABCA1 deficiency affects SREBP2-dependent cholesterol homeostatic responses *in vivo*. We show that SREBP2 response gene expression is partly downregulated in the liver of *Abca1*^-/-^ mice compared to wild-type mouse livers in the steady-state condition. In contrast, we find that in fasted or refed condition, the expression of SREBP2 target genes is upregulated in multiple tissues of *Abca1*^-/-^ mice. Correspondently, SREBP2 processing is also increased in select tissues of these mice. Altogether, our results suggest that ABCA1 deficiency is associated with the tissue-specific dysregulation of the SREBP2 pathway in a manner dependent on nutritional status. Lack of ABCA1 thus causes not only HDL deficiency but also dysregulation of the SREBP2 pathway in multiple tissues, which may underlie the pathophysiology of TD.

## Introduction

High-density lipoprotein (HDL) levels are inversely correlated to the incidence of cardiovascular disease (CVD), and HDL exerts many beneficial effects on preventing cardiovascular events, including reverse cholesterol transport (RCT) and anti-inflammatory and anti-oxidative properties [1-4]. The ATP-binding cassette transporter ABCA1 plays an indispensable role in the formation of HDL. Mechanistically, it exports cellular phospholipid and cholesterol to lipid-free apolipoprotein A-I (apoA-I), a major apolipoprotein that constitutes HDL, from the plasma membrane (PM), which results in the formation of nascent HDL [5,6]. In humans, loss-of-function mutations in the *ABCA1* gene cause Tangier disease (TD), a familial HDL deficiency [7-9]. Despite reduced low-density lipoprotein (LDL)-cholesterol levels, TD patients often develop premature atherosclerosis and have a higher risk of CVD [10,11]. In addition, rare mutant alleles in this gene are associated with low HDL levels in humans [12]. Furthermore, genetic ablation of the *Abca1* gene in mice results in the disappearance of plasma HDL, and these mice are more prone to develop atherosclerotic lesions when challenged with the Western diet [13].

In addition to HDL deficiency, lack of ABCA1 leads to the deposition of cholesterol in peripheral tissues, which is a hallmark of TD pathophysiology [10]. In addition to impaired RCT, it is suggested that altered sterol metabolism is involved in the development of premature atherosclerosis in TD patients. In *Abca1*^-/-^ mice, the cholesterol biosynthesis rate is altered in a tissue-specific manner; in some tissues, including the adrenal, small bowel, and colon, the rate of synthesis is increased, whereas it is decreased in the lung and unchanged in the liver [14]. Other studies have also shown a tissue-specific dysregulation of HMG-CoA reductase (HMGCR) activity, a rate-limiting enzyme of cholesterol biosynthesis; its activity is higher in the adrenal and spleen, but lower in the liver from *Abca1*^-/-^ mice [15]. Furthermore, LDL receptor expression and LDL clearance are markedly increased in the liver of hepatocyte-specific *Abca1*^-/-^ mice [16]. At cellular levels, it has been shown that ABCA1 deletion leads to the activation of sterol regulatory element-binding protein 2 (SREBP2) processing and its target gene expression [17,18]. Moreover, in hepatoma cells, the reduction of ABCA1 expression by p53 mutation activates the SREBP2 pathway, which contributes to the development of hepatocellular carcinoma [18]. On the other hand, none of previous studies has examined SREBP2 processing nor more comprehensive SREBP2 response gene expression in multiple tissues of *Abca1*^-/-^ mice. In addition, although SREBP2 activity is highly regulated by nutritional status in vivo [19], whether lack of ABCA1 alters the activity in different nutritional conditions is unknown. As such, the in vivo significance of ABCA1 deficiency in sterol regulation remains incompletely understood.

To elucidate tissue cholesterol homeostasis in TD, we investigated whether ABCA1 deficiency affects SREBP2 activity in vivo under different nutritional conditions using *Abca1*^-/-^ mice. Our results suggest that ABCA1 deficiency alters the SREBP2-dependent gene expression in tissue-specific and nutritional status-dependent manners in mice.

## Materials and Methods

### Animal Studies

*Abca1*^+/-^ mice (DBA/1-Abca1^tm1Jdm^/J) were purchased from Jackson’s Animal Laboratories. They were backcrossed onto C57BL/6 mice, and heterozygous N3 animals were intercrossed to obtain offspring [20]. Mice were fed *ad libitum* with a standard chow diet unless noted otherwise, maintained in a pathogen-free environment, and kept on a 12-h light/12-h dark schedule. The genotypes of all offspring were determined as described [20]. We used 2–4-month-old male mice. For fasting and refeeding experiments, mice were fasted for 20–24 h, then refed with a normal chow diet for 10–12 h. After mice were fasted or refed, they were euthanized by cervical dislocation. Tissues (liver, adrenal, spleen, and brain (cerebrum)) were removed, immediately frozen in liquid nitrogen, and stored at −80ºC until use. The experimental procedures were pre-approved by the Animal Welfare Committee of Nagoya City University Graduate School of Medical Sciences and according to institutional guidelines.

### Quantitative RT-PCR (RT-qPCR)

Total RNA was isolated with TRIzol Reagent (Thermo Fisher Scientific). mRNA levels of various genes were determined by quantitative real-time PCR and quantified by the ΔΔCT method; *Hprt* expression was used as an internal control as previously described [21]. Primers for each gene are described previously [17,21].

### Immunoblot analysis

Nuclear extracts of mouse liver and spleen were prepared according to a method previously described [22]. Mouse adrenal tissue lysates were prepared by homogenizing (150 μl/adrenal) in urea buffer (8 M urea, 50 mM Tris-HCl pH 8.0, 150 mM NaCl) containing 25 μg/ml ALLN (Wako) and protease inhibitor cocktail (Sigma-Aldrich)[23]. Protein concentration was determined by BCA Protein Assay (Thermo Fisher Scientific). Aliquots of nuclear extracts or tissue lysate were subjected to SDS-PAGE and immunoblot analysis. β-actin and p36/MAT1 were used as loading controls for whole tissue lysate and nuclear extracts, respectively.

Hybridoma cells producing anti-HMG-CoA reductase antibody (IgG-A9) were obtained from ATCC, and the supernatant of hybridoma cell cultured media were used for immunoblotting. Other antibodies were obtained from commercial sources as follows: rabbit anti-SREBP2 polyclonal antibody from Cayman (#10007663), mouse anti-β-actin monoclonal antibody (clone AC15) from Sigma (A5441), and mouse anti-p36/MAT1 monoclonal antibody from BD Biosciences (#610532).

### Statistical Analyses

Data are presented as means ± SD. Statistical analysis was performed using a two-tailed, unpaired Student’s *t*-test. A *p* value less than 0.05 was considered statistically significant.

## Results

### Steady-state expression of SREBP2 response genes in Abca1^-/-^ mouse livers

In addition to HDL formation, ABCA1 is involved in the regulation of SREBP2 at cellular levels [17,18]. We therefore examined whether ABCA1 deficiency alters hepatic SREBP2 activity in vivo in the non-fasted, steady-state condition. We compared the expression of various SREBP2 target genes involved in cholesterol biosynthesis (*Hmgcs1, Hmgcr, Fpps, Sqs, Cyp51a1*) and uptake (*Ldlr*) in the liver of wild-type (WT) and *Abca1*^-/-^ mice fed normal chow diet. The results show that only mRNA levels of *Fpps* (which encodes farnesyl pyrophosphate synthase) are significantly reduced in *Abca1*^-/-^ mice (**Fig. 1**). The hepatic expression of other SREBP2 response genes was not altered between WT and *Abca1*^-/-^ mice, which is largely consistent with a previous report [24]. The results suggest that ABCA1 deficiency shows almost no effects on hepatic SREBP2 response gene expression in the steady-state condition.

**Figure 1.**
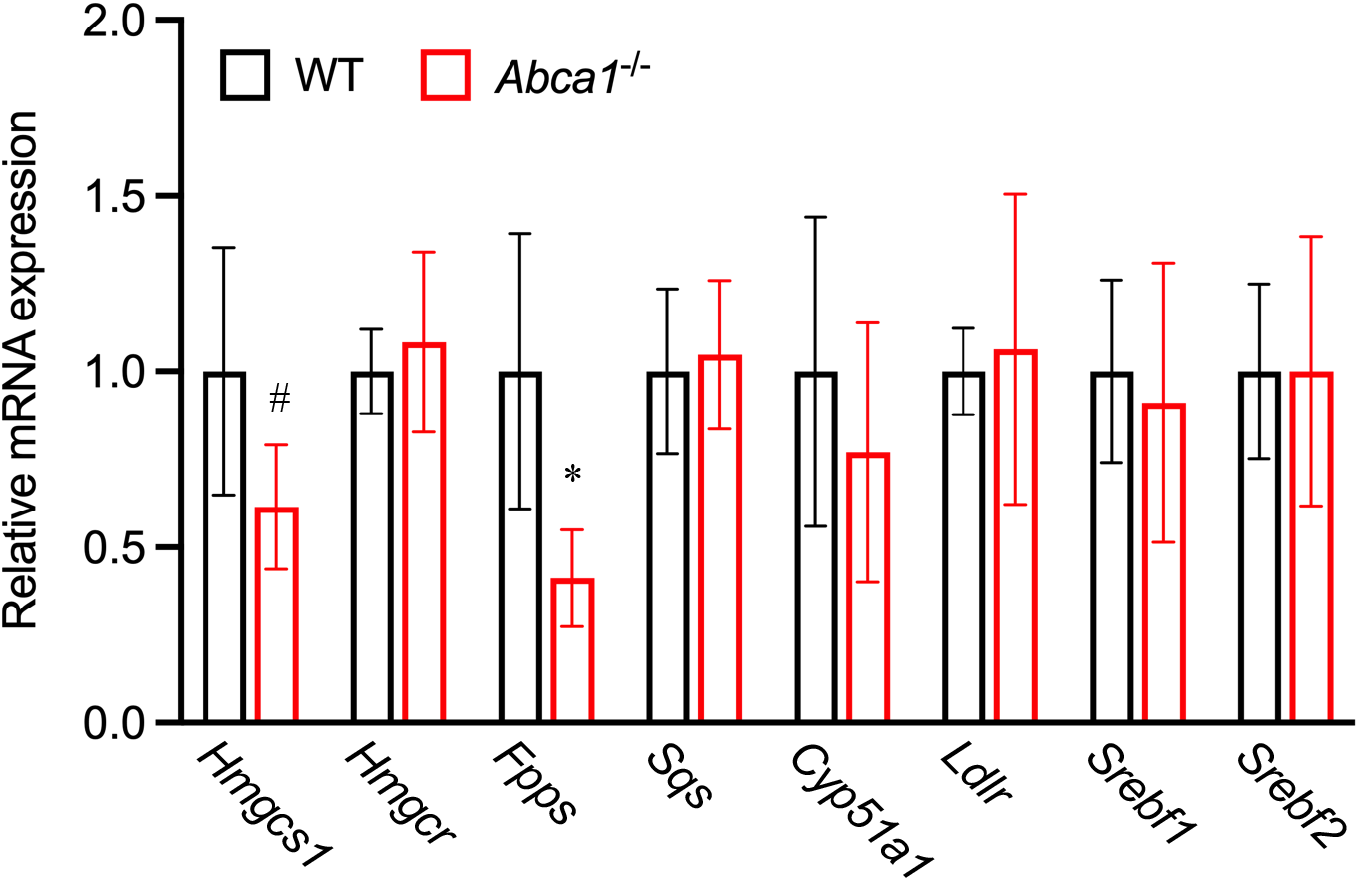
Expression of SREBP2 response genes in livers of *Abca1*^-/-^ mice in a steady-state condition. mRNA levels of SREBP2 response genes in livers from WT and *Abca1*^-/-^ mice fed a normal chow diet were examined by qRT-PCR. Data represent mean ± SD (n=4). Statistical analyses were performed by student’s *t*-test. ^#^ *p* < 0.1, * *p* < 0.05.

### SREBP-2 response gene expression in multiple tissues of WT and Abca1^-/-^ mice under the fasted and refed conditions

The activity of SREBP2 is highly dependent on nutritional status such as fasting and refeeding in the liver [19]. To further explore the effect of ABCA1 deficiency on SREBP2 activity, we examined the expression of various SREBP2 response genes (*Hmgcs1, Hmgcr, Fpps, Sqs, Cyp51a1, Ldlr, Pcsk9*, and *Srebf2*) not only in the liver but also in other tissues, including the adrenal, spleen, and brain (cerebrum) isolated from fasted and refed mice. All these four tissues express significant levels of ABCA1 both at mRNA and protein levels [25]. The age-matched male WT and *Abca1*^-/-^ mice were fasted overnight followed by refeeding with a normal chow diet, based on a previously published protocol [19]. Among the four tissues, the most remarkable differences between WT and *Abca1*^-/-^ mice were observed in the adrenal. Under both fasted and refed conditions, adrenals from the *Abca1*^-/-^ mice exhibit markedly elevated expression of almost all SREBP2 response genes examined (**Fig. 2A**), but *Srebf1* mRNA (which encodes SREBP1) levels were unchanged between WT and *Abca1*^-/-^ mice.

**Figure 2.**
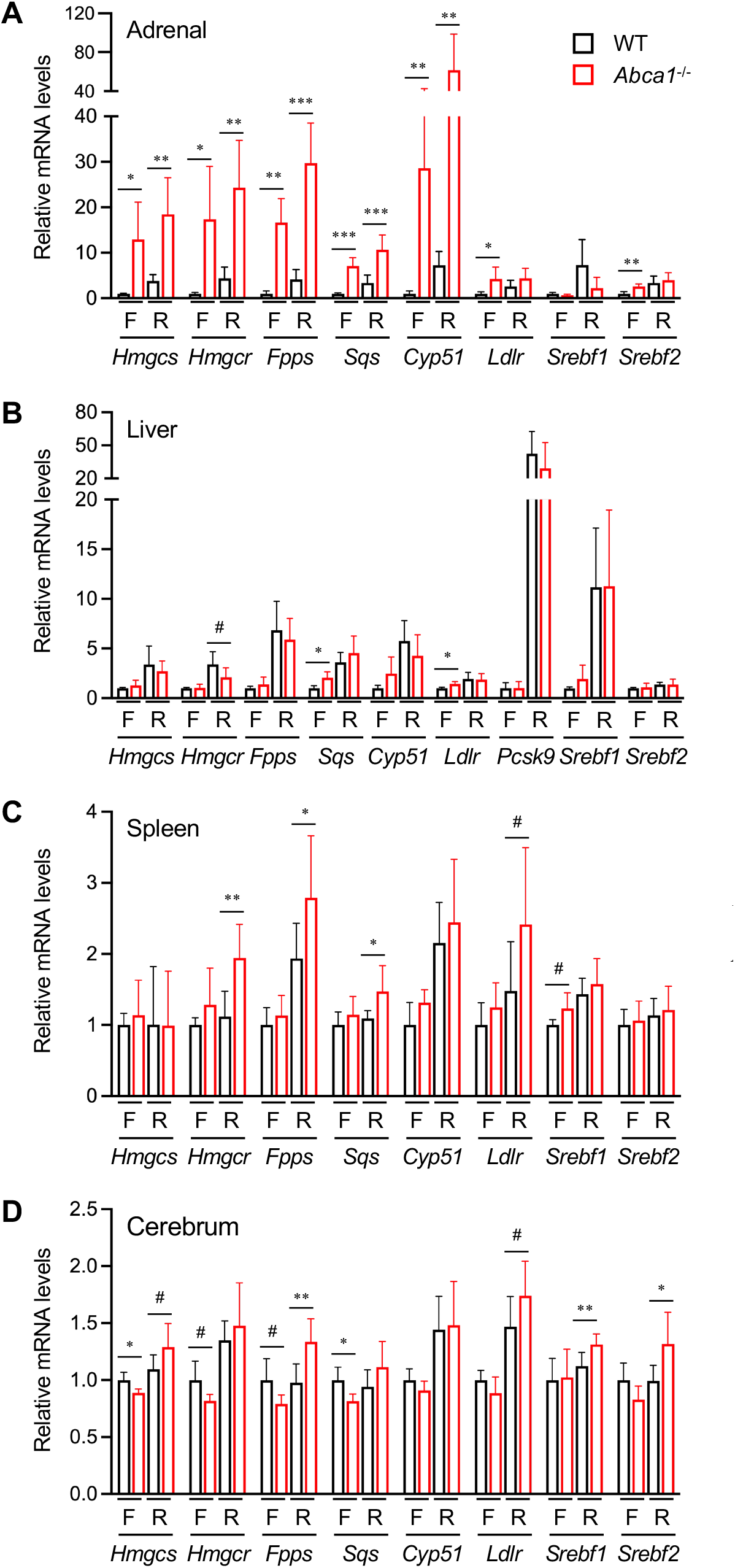
Expression of SREBP2 response genes in multiple tissues of *Abca1*^-/-^ mice under fasted and refed conditions. WT and *Abca1*^-/-^ mice were fasted (F) and refed (R) as described in Materials and Methods. mRNA levels of SREBP2 target genes in the adrenal (A), liver (B), spleen (C), and cerebrum (D) were analyzed by qRT-PCR. The results represent relative changes to WT mice under the fasted condition. Results are mean ± SD (n=4 for the fasted group, n=6–8 for the refed group). Student’s *t*-test was performed between WT and *Abca1*^-/-^ mice in each condition. ^#^ *p* < 0.1, * *p* < 0.05, ** *p* < 0.01, *** *p* < 0.001.

In the liver, in addition to *Srebf1*, refeeding markedly increased the expression of most SREBP2 response genes both in WT and *Abca1*^-/-^ mice (**Fig. 2B**). The results are good agreement with previous studies showing that refeeding activates both SREBP1 and SREBP2 in this tissue [19]. PCSK9 was the most elevated gene among them. Although there were no differences between WT and *Abca1*^-/-^ mice in mRNA levels of all the SREBP2 response genes in the refed condition, small increases in mRNA levels of *Sqs* and *Ldlr* in *Abca1*^-/-^ mice compared to WT mice were observed under the fasted condition. These results suggest that ABCA1 deficiency slightly induces SREBP2 response gene expression in the liver only under the fasted condition.

We next examined the SREBP2 response gene expression in the spleen. In *Abca1*^-/-^ mice under the refeeding condition, elevated expression of several SREBP2 response genes was observed in the spleen although the magnitudes of the increases were modest (**Fig. 2C**). Statistical analysis shows that the differences in mRNA levels of *Hmgcr, fpps*, and *Sqs* between WT and *Abca1*^-/-^ mouse spleens under the refed condition were significant. On the other hand, none of the SREBP2 response genes was altered in *Abca1*^-/-^ mouse spleens under the fasted condition.

Finally, we compared mRNA levels of SREBP2 target genes in the cerebrum between WT and *Abca1*^-/-^ mice. Under the fasted condition, the expression of several SREBP2 target genes (*Hmgcs1, Hmgcr, Fpps*, and *Sqs*) was slightly downregulated in the cerebrum of *Abca1*^-/-^ mice compared to WT mice (**Fig. 2D**). In contrast, under the refed condition, we found the higher mRNA levels of several SREBP2 target genes, including *Hmgcs1, Fpps, Ldlr*, and *Srebf2* in the cerebrum of *Abca1*^-/-^ mice than those in WT mice.

### SREBP2 processing in WT and Abca1^-/-^ mice

The proteolytic cleavage of precursor SREBP2 to generate the active mature form is a highly regulated process for upregulating the expression of the target genes. This cleavage is inhibited by fasting but is activated by refeeding in the liver [19]. To directly demonstrate the activation of SREBP2, we examined the processing in the adrenal, liver, and spleen where increases in SREBP2 response gene expression were observed in *Abca1*^-/-^ mice. SREBP mature form was undetectable in the cerebrum although nuclear extracts were prepared using essentially the same protocol with the liver and spleen. We prepared whole cell extracts from the adrenal (because the adrenal is much smaller than the liver and spleen) or nuclear extracts from the liver and spleen of WT and *Abca1*^-/-^ mice and assessed precursor and mature forms in the adrenal and mature form in the spleen and liver by immunoblot. The results show that in the adrenal of *Abca1*^-/-^ mice under both fasted and refed conditions, the SREBP2 mature form was markedly elevated (**Fig. 3A**), which is in line with mRNA analyses shown in Figure 2. HMGCR protein expression was also increased in *Abca1*^-/-^ mouse adrenals under both the fasted and refed conditions. Similar to the recent findings in the liver [26], refeeding further induced HMGCR protein expression compared to the fasted condition in the adrenal of *Abca1*^-/-^ mice. In WT mouse adrenals, HMGCR protein expression was as low as undetectable levels. We next examined the mature form of SREBP2 in the liver. Consistent with the induction of most SREBP2 response gene expression, refeeding increased the mature form in the liver of WT and *Abca1*^-/-^ mice (**Fig. 3B**). Only under the fasted condition, the mature form was increased by 1.7-fold in the liver of *Abca1*^-/-^ mice compared to that of WT mice. These results are largely consistent with mRNA expression analysis (**Fig. 2B**). We also found that in the spleen, *Abca1*^-/-^ mice showed 2- to 3-fold increases in SREBP2 mature form not only in refed condition but also in the fasted condition (**Fig. 3C**). Together, levels of SREBP2 mature form in these three tissues may explain the increases in mRNA levels of SREBP2 response genes in *Abca1*^-/-^ mice.

**Figure 3.**
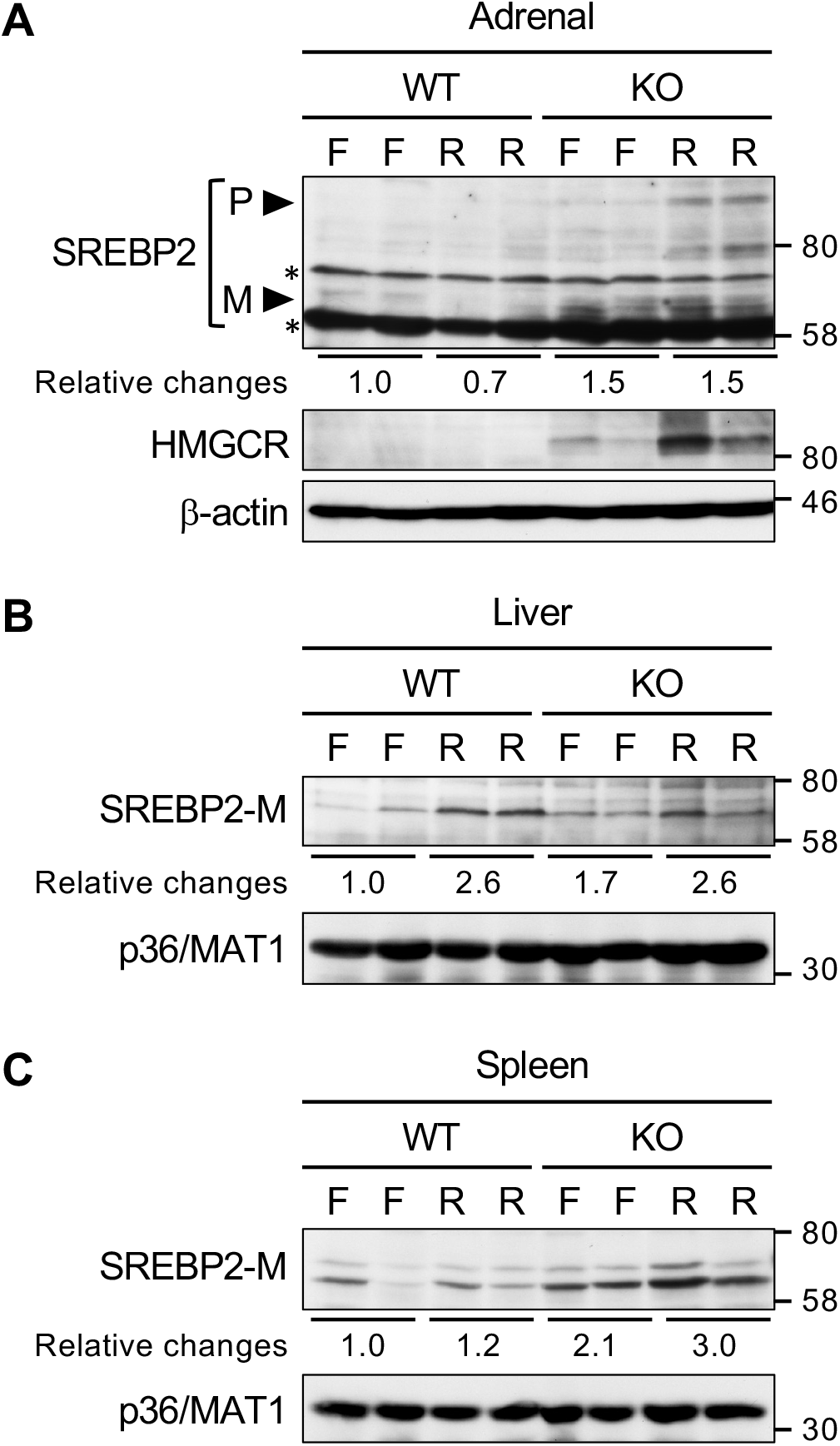
SREBP2 processing in multiple tissues of *Abca1*^-/-^ mice. Whole tissue extracts of the adrenal (A) or nuclear extracts of the liver (B) and spleen (C) were prepared from two mice in the fasted (F) and refed (R) conditions for examining SREBP2 processing and HMGCR expression (adrenal only). Equal amounts of protein were subjected to immunoblot analysis to detect SREBP2 and HMGCR proteins. β-actin or p36/MAT1, a nuclear protein, was used as a loading control. Relative changes in the SREBP2 mature form shown below the SREBP2 immunoblots are the average values of the two mice. Asterisks denote non-specific bands. P, precursor form; M, mature form.

## Discussions

Studies with tissue-specific *Abca1* knockout mice revealed that the liver produces 70– 80% of plasma HDL [27] whereas the rest (20–30%) of HDL is produced in the intestine [28]. On the other hand, ABCA1 is expressed not only in these two tissues but also in other tissues [25], suggesting that it exerts tissue-specific roles. Cholesterol deposition in tissues is a hallmark of TD pathophysiology [10], yet the mechanism has not been fully addressed. Studies with various cultured cells showed that ABCA1 regulates membrane structures and functions, including transbilayer cholesterol distribution in the PM, cholesterol contents in lipid rafts, clathrin-independent endocytosis, and membrane curvature formations [29,30]. In addition, we and others have shown that lack of ABCA1 induces SREBP2 processing by limiting endoplasmic reticulum cholesterol, which upregulates the expression of various SREBP2 response genes involved in cholesterol biosynthesis and uptake [17,18]. However, the role of ABCA1 in the regulation of SREBP2 in vivo has been incompletely elucidated.

Our current results suggest that ABCA1 deficiency alters SREBP2 activity in tissue-specific and nutritional status-dependent manners in multiple tissues. The abnormalities in cholesterol homeostasis were manifest most severely in the adrenal, less severely in the spleen and brain, and least severely in the liver in *Abca1*^-/-^ mice. We showed that in the adrenal and spleen of *Abca1*^-/-^ mice, expression of SREBP2 response genes and SREBP2 processing are upregulated in the fasted and/or refed conditions while the degree of the induction was much higher in the adrenal. These results are largely consistent with previous observations reporting the increases in cholesterol synthesis, HMGCR activity, and LDL uptake in the adrenal and spleen of *Abca1*^-/-^ mice [14,15]. Thus, our results demonstrate that the changes observed in these tissues are associated with elevated SREBP2 activity. The differences in the magnitude of the increases could be explained by a tissue-dependent cholesterol demand; adrenals require more cholesterol to synthesize steroid hormones, thereby upregulating the SREBP2 pathway to compensate for the lack of HDL. We also found that only in the fasted liver, a few SREBP2 response genes exhibit slightly higher expression in *Abca1*^-/-^ mice than in WT mice. No previous studies with *Abca1*^-/-^ mice were examined and compared the SREBP2 pathway between fasted and refed conditions.

Refeeding can boost the expression of not only SREBP1c target genes but also SREBP2 response genes in the liver [19]. In addition to transcriptional regulation, refeeding stabilizes HMGCR protein by activating the mTORC1-USP20 axis in the liver, contributing to the further promotion of cholesterol biosynthesis [26]. In mice with hepatocyte-specific ABCA1 deficiency, a partial impairment in liver insulin signaling and SREBP1c-dependent lipogenesis has been reported [31], but whether the SREBP2 pathway is also affected in these mice is unknown. Our results show that the induction of SREBP2 processing and its target gene expression is comparable under the refed condition between WT and *Abca1*^-/-^ mouse livers, suggesting that signaling events for activating the SREBP2 pathway are not impaired at least in the liver of global *Abca1*^-/-^ mice. Furthermore, it is of interest that we observed opposite effects of ABCA1 deficiency on the expression of SREBP2 response genes in the brain (cerebrum) between the fasted and refed conditions although the differences are only subtle; in the fasted condition, SREBP2 response gene expression was reduced whereas the expression was upregulated in the refed condition in *Abca1*^-/-^ mice. A mechanistic explanation for these changes is difficult at present. However, it has been reported that intermediate sterols such as lathosterol and desmosterol are increased in the cortex and hippocampus of brain-specific *Abca1* knockout mice, suggesting that ABCA1 deficiency increases cholesterol biosynthesis in the brain [32]. The results may explain our findings that the expression of several cholesterol biosynthetic genes is upregulated in the cerebrum of *Abca1*^-/-^ mice under the refed condition. Since ABCA1 and cholesterol homeostasis play diverse important roles in neurological diseases, including Alzheimer’s disease [33], further studies on the regulation of brain cholesterol homeostasis are required to reveal a mechanistic link between cholesterol metabolism and complex neurological diseases.

In summary, the present study shows that ABCA1 deficiency is associated with dysregulation of the SREBP2 pathway in multiple tissues in fasted or refed conditions. Our findings may have a clinical significance; the upregulation of SREBP2 activity by ABCA1 deficiency could partly underlie the cholesterol deposition in the peripheral tissues of TD patients.

## Acknowledgements

We thank Drs. Ta-Yuan Chang and Maximillian Rogers (Geisel School of Medicine at Dartmouth) for discussion, and Haruka Hayashi for mouse genotyping.

